# Single-cell mutation rate of turnip crinkle virus (-)-strand replication intermediates

**DOI:** 10.1101/2023.05.03.539190

**Authors:** Camila Perdoncini Carvalho, Junping Han, Khwannarin Khemson, Ruifan Ren, Luis Eduardo Aranha Camargo, Shuhei Miyashita, Feng Qu

## Abstract

Viruses with single-stranded, positive-sense (+) RNA genomes incur high numbers of errors during replication, thereby creating diversified genome populations from which new, better adapted viral variants can emerge. However, a definitive error rate is known for a relatively few (+) RNA plant viruses, due to challenges to account for perturbations caused by natural selection and/or experimental set-ups. To address these challenges, we developed a new approach that exclusively profiled errors in the (-)-strand replication intermediates of turnip crinkle virus (TCV), in singly infected cells. A series of controls and safeguards were devised to ensure errors inherent to the experimental process were accounted for. This approach permitted the estimation of a TCV error rate of 8.47 X 10^−5^ substitution per nucleotide site per cell infection. Importantly, the characteristic error distribution pattern among the 50 copies of 2,363-base-pair cDNA fragments predicted that nearly all TCV (-) strands were products of one replication cycle per cell. Furthermore, some of the errors probably elevated error frequencies by lowering the fidelity of TCV RNA-dependent RNA polymerase, and/or permitting occasional re-replication of progeny genomes. In summary, by profiling errors in TCV (-)-strand intermediates incurred during replication in single cells, this study provided strong support for a stamping machine mode of replication employed by a (+) RNA virus.

**Author Summary:** Most (+) RNA viruses introduce replication errors at relatively high frequencies. As a result, it is of vital importance for these viruses to purge lethal errors in a timely manner. TCV, a plant-infecting small (+) RNA virus, was proposed to encode a Bottleneck, Isolate, Amplify, Select (BIAS) mechanism that compel swift clearance of lethal errors by bottlenecking the number of replicating genome copies to one per cell. A crucial prediction of this BIAS model is that such bottlenecking also acts on progeny genome copies, preventing them from repeating replication in the cells of their own genesis. The current study tested this prediction by developing a carefully controlled, readily reproducible approach to profile errors and error distributions in (-)-stranded replication intermediates of TCV. We found that most of replication-generated (-) strands descended from the primary (+) strands through a single replication cycle. This finding adds fresh support to the BIAS model.

## Introduction

Viruses with single-stranded, positive sense (+) RNA genomes are a major class of human, animal, and plant pathogens that include poliovirus, Dengue virus, Zika virus, and more recently SARS-CoV-2 causing the global COVID-19 pandemic. These viruses are known to incur high numbers of errors during the process of genome replication, as the virus-encoded RNA-dependent RNA polymerases (RdRps) mostly lack proofreading activities (1–4). While many of the replication errors threaten viral competitiveness or even viability, they also create reservoirs of diversity through which new variants can emerge. Some of the new variants may enable a virus to spread and flourish in a species different from its original host, causing devastating diseases in the new host species. Therefore, knowledge about the mutation rates of (+) RNA viruses is critical for assessing the potential risks of emerging viruses, especially those that transcended the host barriers in the recent past.

However, mutation rates are known for a surprisingly few (+) RNA viruses, primarily due to difficulties to account for biases caused by natural selection, and/or the measurement process itself. Several frequently used methods have been critically reviewed by Peck and Lauring (2). The most common ones are Sänger sequencing of randomly selected cDNA clones of viral genomes produced during infections of culture cells or host individuals (5–8), and more recently, high-throughput sequencing of viral cDNA libraries (9, 10). Despite various precautions taken to minimize selection biases or mutations introduced by the measurement procedures such as reverse transcription-polymerase chain reaction (RT-PCR), mutations identified by these two methods likely still favor more competitive ones, while undercounting deleterious ones. A third approach known as mutation accumulation assay uses serial dilutions to separate genome variants from each other, subsequent parallel propagation of these variants permits the capture of variants less capable of competing in a mixed infection environment. The least biased method is the fluctuation assay, which relies on a scorable, selection-neutral phenotype that can be abolished, and restored, with very few point mutations. Spontaneous reversions or compensatory mutations occurring during virus replication that restore the abolished phenotype are then profiled through phenotype scoring and/or sequencing (11, 12). A drawback of this approach is that huge numbers of phenotype-restoring variants must be screened in order to capture the complete spectrum of mutations necessary for robust mutation rate computation. This is because the initial phenotype-abolishing change(s) usually alter just one or very few nucleotides of a viral genome. Accordingly, the chance of spontaneous phenotype-restoring mutations occurring at these few sites would be extremely low, necessitating the screening of enormous numbers of descendant copies. Finally, most of these methods required a pre-propagation step to bulk up the inoculum pools. As a result, mutations occurring during this pre-propagation stage may distort the estimation of mutation rates (11, 13).

To address these limitations, we initiated a study to obtain a more accurate estimation of mutation rates incurred during replication of a (+) RNA virus. We decided to focus our attention on (-)-strand RNA intermediates [referred to as (-) strands hereafter for simplicity] produced during (+) RNA virus replication in single cells. This decision was based on the understanding that production of (-) strands was the first step of (+) RNA virus replication, thus errors detectable in the (-) strands in primarily infected cells would directly reflect the mutation rate of a viral RdRp, commonly defined as substitutions per nucleotide site per cell infection (s/n/c) (2). Therefore, the hypothesis for the current study was that (-) strands in single cells, having been exposed to minimal selection pressure, retained the full spectrum of replication errors, including lethal ones.

An added benefit of analyzing (-) strands was to permit the estimation of viral replication cycles per cell. This is because, should (-) strands be produced from the (+) strands of progeny viruses arisen from repeated cycles of replication, the number of errors in them would increase at a rate of two folds for every additional cycle [(-) strand to (+) strand, then back to (-) strand]. This would cause cells to contain a mixture of (-) strands containing varying number of errors depending on the number of replication cycles they repeated. As a result, individual (-) strands sampled for sequencing would be expected to contain errors whose numbers varied dramatically from each other, and severely deviate from the error frequency mean obtained by dividing the total number of (-) strands with the total number of errors. More precisely, they would deviate from the random distribution pattern governed by the Poisson distribution law. In short, by comparing the observed error distributions in different (-) strands with Poisson distribution predictions, we would be able to determine whether all (-) strands in an average cell descended from a single primary (+) strand through a stamping machine replication mode. The Poisson distribution rationale was invoked by previous authors (13) to conclude that bacteriophage ɸ6 with a double-stranded RNA genome replicated through a predominantly stamping machine mode.

To test the feasibility of our idea, we adopted turnip crinkle virus (TCV) as the model for the current study. TCV is a small virus belonging to the Genus *Betacarmovirus*, Family *Tombusviridae*, with a (+) RNA genome of 4,054 nucleotides (nt) (Fig. 1). TCV genomic RNA (gRNA) encodes five proteins: the 5’ proximal p28 and its translational read-through product p88 are both essential for TCV replication. Two subgenomic RNAs (sgRNA1 and 2) are produced inside the infected cells, with sgRNA1 serving as mRNAs for the p8 and p9 movement proteins (MPs) (14, 15), and sgRNA2 the p38. p38 is both the viral capsid protein (CP), and the TCV-encoded suppressor of RNA silencing (VSR) (16–18). Our results suggested that TCV RdRp incurred errors at a rate of 8.47 X 10^−5^ for every nucleotide incorporated in the (-) strands in single cells. This rate translates into approximately 0.69 error for every new (+) TCV genome synthesized, and is within the range of previous estimates obtained with other plant-infecting (+) RNA viruses (2, 19). Notably, our approach permitted detection of errors with lethal consequences. Finally, the observed pattern of error distribution in individual (-) strands suggested that progeny TCV genome copies rarely repeat replication in the cells of their own genesis.

**Figure 1.**
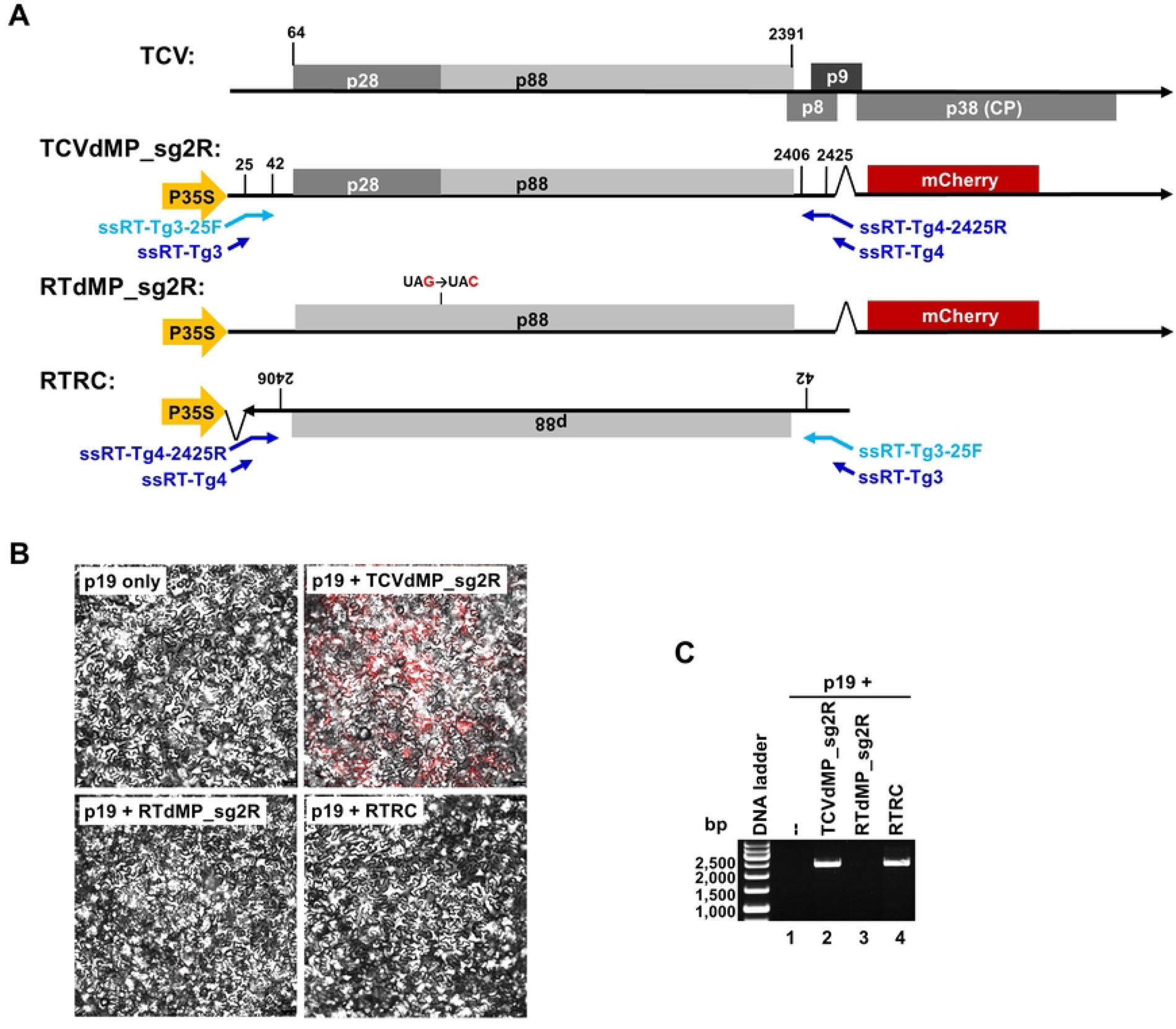
Generation of (-)-strand-specific RT-PCR products of TCV for error profiling. **A**. Constructs assembled for the current study. The top diagram depicts the (+)-strand genome of TCV encoding proteins p28, p88, p8, p9, and p38. Among them, p8 and p9 are translated from sgRNA1, p38 sgRNA2. TCVdMP_sg2R is a modified TCV replicon integrated in the binary plasmid pAI101, downstream of the strong P35S promoter so that once inside plant cells, the host cell PolII would be recruited to transcribe replication-initiating viral RNAs. Note that the p8 and p9 MP ORFs were deleted, and the p38 ORF was replaced by that of mCherry. Note that the short, arrowed lines in blue symbolize the four primers used in strand-specific RT-PCR, with the one in light blue used in the RT step, whereas the three in dark blue used in the PCR step. RTdMP_sg2R was further modified from TCVdMP_sg2R by eliminating the p28 stop codon, rendering its transcripts incapable of launching replication, hence are exclusively (+) strands. RTRC contains the reverse-complemented form of the first 2,489 nt of RTdMP_sg2R, so that P35S-driven transcription leads to RNAs that are exclusively (-)-stranded relative to TCV genome. **B**. Confocal microscopy images of N. benthamiana epidermal cells showing that only TCVdMP_sg2R replicated to produce mCherry fluorescene. **C**. Strand-specific RT-PCR showing that only the two constructs expected to synthesize (-) strands, TCVdMP_sg2R and RTRC, produced PCR products of expected size (2,363 bp). Note that at 24 PCR cycles, the RTRC-specific product was much less abundant. D. Serial dilution, followed by RT-PCR, showing TCVdMP_sg2R-derived (-)-strand cDNA was approximately 9 times more abundant than that of RTRC.

## Results and Discussion

### Strategy to capture errors in the (-) strands of replicating TCV with high confidence

In an effort to determine the error rate of TCV RdRp, we introduced several measures to overcome various limitations associated with existing error-profiling methods (Table 1). First, to address the uncertainty related to errors introduced into viral inoculums through pre-propagation, we delivered TCV cDNA directly into plant cells to initiate viral infections. To accommodate such delivery, the full-length TCV cDNA was inserted in a binary plasmid (pAI101) (20–22), under control of the strong 35S promoter (P35S) of cauliflower mosaic virus (CaMV). The resulting plasmid was sent into cells of *Nicotiana benthamiana* plants via agro-infiltration (see Materials and Methods for details). While the viral cDNA still needed to be transcribed into replication-initiating RNA upon entering plant cells, we reasoned that the DNA-dependent RNA polymerase II (Pol II) of *N. benthamiana* cells, recruited by P35S to carry out transcription, would incur relatively few errors (23, 24). Although this assumption was proven to be inaccurate by our data, Pol II-introduced errors were easily controlled with a non-replicating construct (RTRC. Fig 1A, bottom) that produced (-) strands only, in a Pol II-dependent manner (see later).

**Table 1.** Experimental design – challenges and solutions

| Challenge | Solution |
| --- | --- |
| 1. Replication-initiating viruses might contain variable numbers of founding errors | Initiate replication with RNA transcribed by host Pol II that incurs one round of errors at a constant rate. |
| 2. Viral genomes might have replicated in a variable number of cells prior to sampling | Restrict replication in single cells by using a plant-infecting virus (TCV) with MP genes disrupted. |
| 3. Phenotype-based selection might enrich certain mutations | Profile errors in the non-protein-coding (-)-strand replication intermediates. |
| 4. Differentiate between primary and secondary (-) strands | Generate and analyze continuous cDNA fragments of at least 2,000 nt in length. |
| 5. Ensure cDNA of (-) strands is exclusively derived from (-) strands | a. Carry out strand-specific RT-PCR following an established procedure;<br>b. Include control constructs that produce exclusively (+) and (-) strands, respectively. |
| 6. Account for errors incurred during all steps of the error-profiling experiment | Include appropriate controls at every step. Especially relevant to current study was the RTRC control producing (-) strands independent of viral replication. |

Second, to avoid errors introduced sequentially through reiterative replication in successive cells, TCV cDNA was modified to disrupt the MP-coding region, thereby restricting viral replication in single cells. Additionally, to permit microscopical monitoring of viral replication, the TCV cDNA was further modified to encode an mCherry reporter in place of p38, enabling replication-dependent expression of the mCherry fluorescent protein. Previous studies (17, 25) established that TCV replication in single cells was not compromised by loss of p38, as long as its VSR function was compensated by a transiently expressed VSR of a different virus, which in our experiments was p19 of tomato bushy stunt virus (TBSV) (16, 25). Combining these modifications led to the TCVdMP_sg2R replicon (Fig 1A & B).

Most importantly, to preclude natural selection-based differential amplification of mutation-containing genomes, we focused on errors incurred in the process of synthesizing viral (-) strands. We reasoned that should TCV replicate for a single cycle in each cell, all (-) strands would incur similar numbers of errors. On the other hand, were multiple cycles of replication to occur in each cell, such occurrences could be detected in the form of (-) strands containing varying numbers of errors concomitant with the number of replication cycles they experienced. More specifically, using primary (-) strands as reference, error frequencies in 2^nd^, 3^rd^, and 4^th^ generation (-) strands would increase by 3, 5, and 7 folds, respectively, because they would have experienced (-)-to-(+), then back to (-), for one, two, and three rounds, respectively. Therefore, should multi-cycle replication be the predominant mode of replication, we would expect to detect a mixture of (-) strands with dramatically different error numbers.

We further recognized that, given the rarity of errors in short reads, definitive differentiation between these two error distribution patterns would only be possible if the sequence reads were of sufficient lengths (Table 1). Thus, (-)-strand-specific of cDNAs of 2,450 bp in size were produced. These cDNAs were cloned, and individual clones were subject to Sänger sequencing to resolve sequences of the cDNA inserts. Excluding terminal sequences that were part of RT-PCR primers, the sequence usable for error screening was 2,363 bp. This region corresponded to TCV positions 43 – 2,405, encompassing the entire p88 open reading frame (ORF; positions 64 to 2,391. Fig 1 & 2), and accounting for 58% of the 4,054-nt TCV genome.

**Figure 2.**
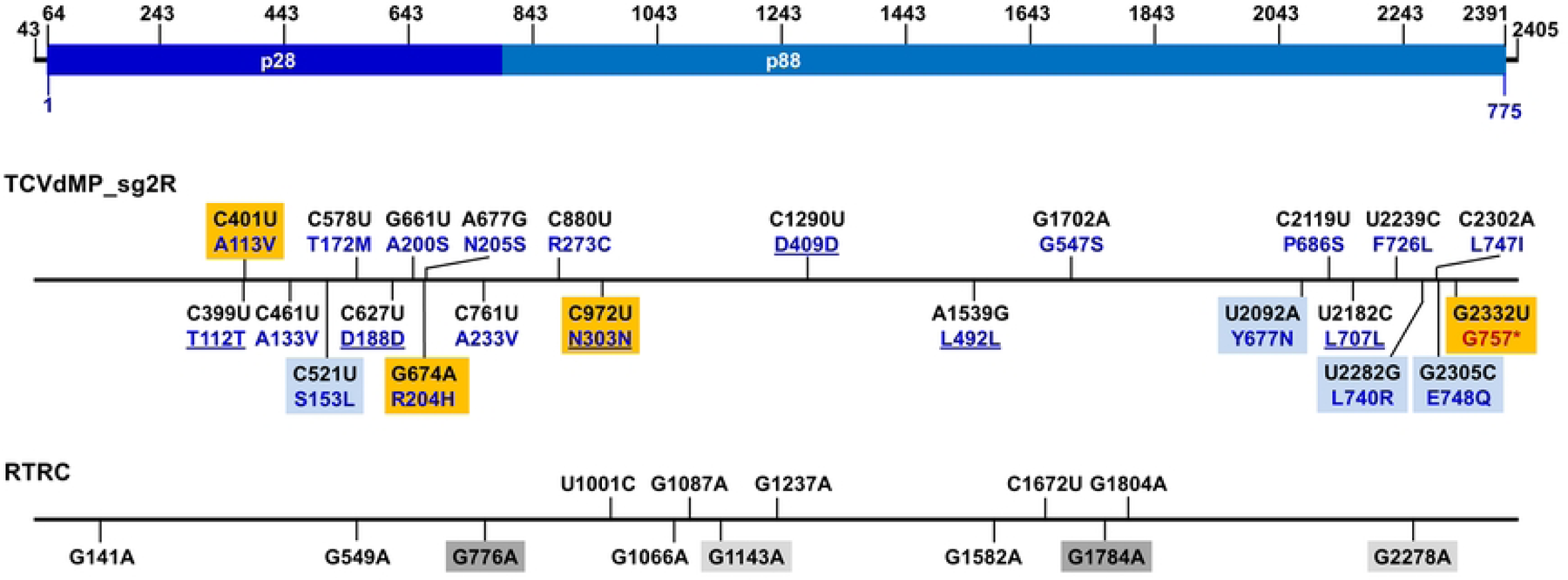
Mapping the identified errors to TCV genome. The top diagram depicts the 2,363-bp range, corresponding to TCV genome positions 43 to 2,435, used for error identification the numbers on top were added as references for convenient allocation of the errors. Within this range, positions 64-2391 encompasses the 775-aa p88 ORF. The middle diagram depicts the 23 errors identified in TCVdMP_sg2R-derived clones, and their corresponding aa changes. The nt changes, converted to (+) polarity, are shown in black font, whereas the corresponding aa changes are shown in dark blue. The unaltered aa residues are underlined. Errors identified from the same clone are highlighted with same-colored boxes (light blue or orange). The bottom diagram depicts 13 errors identified in RTRC-derived clones. Errors occurring in the same clone are highlighted with either light or dark gray boxes.

Another advantage of exclusive profiling of (-) strands was that it permitted unambiguous separation of replication progeny from Pol II transcripts. This is because, while (+) strands in the cell could be Pol II transcripts or progeny of viral replication, (-) strands could only be produced through viral replication. This is important because our previous studies showed that very few (as few as one) of Pol II-transcribed (+) strands initiated TCV replication in each cell, even though thousands of them were present (26–29). Note although (-)-strands with virus sequences could also be produced by host-encoded RNA-dependent RNA polymerases (RDRs), these were normally subject to swift processing to produce small interfering RNAs (siRNAs) (30–32), hence undetectable in sizes longer than 200 nt.

Experimentally, to ensure exclusive use of (-) strands for cDNA synthesis, a strand-specific RT-PCR procedure developed by Plaskon and colleagues (33) was adopted with minor modifications (Fig 1A). Briefly, (-)-strand-specific RT (ssRT) was primed with ssRT-Tg3-25F (Fig 1A, light blue letters and arrows), a chimeric primer containing a 24-nt non-TCV 5’ tail, and an 18-nt TCV-specific 3’ terminus (TCV positions 25-42) complementary to the 3’ end of TCV (-) strands. At the PCR step, ssRT-Tg3, a different primer whose sequence was identical to the non-TCV tail of ssRT-Tg3-25F, was paired with a two-primer mix consisting of ssRT-Tg4 and ssRT-Tg4-2425R, at a 1: 0.05 ratio. ssRT-Tg4 did not contain any TCV sequence, whereas ssRT-Tg4-2425R had the sequence of ssRT-Tg4 as its 5’ tail, and a 20-nt sequence complementary to TCV positions 2,425-2,406 at its 3’ terminus (Fig 1A).

Finally, to control for occasional contamination of (+) strands, and to account for errors incurred during various steps of the experiment, such as Pol II transcription, RT-PCR, cloning, and sequencing, two control constructs were always delivered into plant cells in parallel to TCVdMP_sg2R (Fig 1A). The first, RTdMP_sg2R, contained a 1-nt mutation that abolished the stop codon of p28, causing its transcripts to lose the ability to replicate, thus remaining exclusively (+) sense (Fig 1A). Conversely, the second control, RTRC, harbored the first 2,489 nt of RTdMP_sg2R (including the 1-nt replication-abolishing mutation), but in the reverse-complemented orientation, causing its transcripts to be exclusively (-)-sense (Fig 1A. Also note the landing positions of RT-PCR primers in TCVdMP_sg2R and RTRC). Combining these considerations (Table 1) led to an experimental procedure that yielded highly specific cDNAs of approximately 2,500 bp from RNA samples of TCVdMP_sg2R and RTRC (Fig 1C, lanes 2 and 4), but not those of p19 only or RTdMP_sg2R (lanes 1 and 3) (Also see Fig 1B). These results indicated that the PCR products were derived exclusively from TCV (-) strands.

### Error rate of (-) strands produced by TCV replication

To determine the error rate of TCV RdRp, we then cloned the 2,450-bp (-)-strand-specific PCR products, derived from both TCVdMP_sg2R and RTRC, into the plasmid pUC19. For each of the two PCR products, 50 clones were subject to Sänger sequencing to identify errors. Surprisingly, while 17 out of 50 (34%) clones derived from replicating TCVdMP_sg2R contained one or more errors, 11 out of 50 (22%) clones derived from the non-replicating RTRC also contained errors, making the difference less than two-fold (Table 2). Out of the 17 TCVdMP_sg2R-derived, error-containing clones, 15 had just 1 error each, none had 2 or 3 errors, yet two contained 4 errors each (Table 2). By contrast, among the 11 RTRC-derived, error-containing clones, nine had 1 error each, two had 2 each. The difference in total numbers of errors, 23 versus 13, translated into error rates of 1.947 X 10^−4^ and 1.100 X 10^−4^ s/n/c, for TCVdMP_sg2R and RTRC (-) strands, respectively. An error rate for TCV RdRp was deduced by subtracting the latter rate from the former, resulting in 0.847 X 10^−4^, or 8.47 X 10^−5^. Considering the size of TCV genome (4,054 nt), this error rate meant approximately 34% of (-)-strands synthesized, or 68% of (+)-stranded progeny TCV genome copies, would contain one error, if all progeny genome copies arose from a single cycle of replication. This error rate was within the range of the mutation rates of other (+) RNA plant viruses (5, 8, 19).

**Table 2.** Number of errors in replication-generated (-)-strands of TCVdMP_sg2R, and replication-independent (-)-strands of RTRC

| Construct | TCVdMP_sg2R | RTRC |
| --- | --- | --- |
| Clones sequenced | 50 | 50 |
| Error-free clones | 33 | 39 |
| Error-containing clones | 17 | 11 |
| Clones with 1 error | 15 | 9 |
| Clones with 2 errors | 0 | 2 |
| Clones with 3 errors | 0 | 0 |
| Clones with 4 errors | 2 | 0 |
| Total number of errors | 23 | 13 |
| nt resolved per clone | 2,363 | 2,363 |
| Total nt sequenced | 118,150 | 118,150 |
| Mutation rate (s/n/c) | $1.947 \times 10^{-4}$ | $1.100 \times 10^{-4}$ |

### Unexpectedly frequent error introduction by Pol II

To determine whether certain class(s) of mutations occurred more frequently than others, we next attempted to categorize the specific error classes found in the sequenced DNA fragments. Note that the mutation classes compiled in Table 3 were formatted as if they all occurred in the TCV (-) strands, even though some errors must have been introduced through processes such as Pol II transcription or RT-PCR. As shown in Table 3, the overwhelming majority (11 of 13) of errors derived from the non-replicating (-) strands of RTRC were C-to-U, with the remaining two being A-to-G and G-to-A. Thus, all of RTRC-specific errors were transitions.

**Table 3.** Types of errors [Based on TCV (-)-strands. Stars (*) highlight the fact that the A-to-U, C-to-G, and A-to-C errors occurred in the same clone]

| Error Type |  | TCVdMP_sg2R | RTRC |
| --- | --- | --- | --- |
| Transversion | A → U | 1* | 0 |
|  | U → A | 0 | 0 |
|  | C → G | 1* | 0 |
|  | G → C | 0 | 0 |
|  | A → C | 1* | 0 |
|  | U → G | 0 | 0 |
|  | C → A | 2 | 0 |
|  | G → U | 1 | 0 |
| Transition | A → G | 2 | 1 |
|  | U → C | 2 | 0 |
|  | C → U | 2 | 11 |
|  | G → A | 11 | 1 |
| Indel |  | 0 | 0 |

Previous studies suggested that C-to-U errors, which would be G-to-A in the reverse-transcribed cDNA, were over-represented in reverse-transcriptase-generated errors (23, 24). However, this could not explain why the same class of errors did not occupy a similar percentage of TCVdMP_sg2R-specific clones (2/50 as opposed to 11/50). On the other hand, over-representation of C-to-U errors in RTRC (-) strands could not be attributed to PCR either, as PCR-introduced errors would be expected to have a similar chance to occur in both strands of the double-stranded DNA, hence would manifest as C-to-U and G-to-A with similar frequencies.

However, comparing the C-to-U dominance in RTRC-derived clones (11/50) with the G-to-A dominance in TCVdMP_sg2R-derived clones (11/50) prompted an intriguing revelation. Keep in mind that the RTRC (-) strands were direct transcripts of Pol II, whereas TCVdMP_sg2R (-) strands were copied from (+) stands that were in turn transcripts of Pol II. Therefore, should Pol II be responsible for the C-to-U errors in the RTRC (-) strands, the same class of errors would have been introduced in the TCVdMP_sg2R (+) strands, hence manifesting themselves as G-to-A in TCVdMP_sg2R (-) strands. The fact that these two types of errors occurred in their corresponding (-) strand pools at identical frequencies (both 11/50) was consistent with this interpretation. Thus, Pol II of *N. benthamiana* likely contributed a substantial fraction of errors detected in both (-) strand samples. Conversely, other steps of the experimental procedure, such as RT-PCR and cloning, likely had very modest contributions.

Interestingly, the C-to-U errors were also found to be the most common class of errors introduced by Pol II of *C. elegans* and budding yeast (23, 24). Nevertheless, the error rate of *N. benthamiana* Pol II, at approximately 9.31 X 10^−5^ (11/118,150), was substantially higher than Pol II of *C. elegans* (4 X 10^−6^) and budding yeast (3.9 X 10^−6^) (23, 24). However, in our experiments the *N. benthamiana* Pol II was driven by P35S, a promoter of virus origin. Though not yet investigated, it is possible that Pol II error rate could be affected by the origin and/or strength of the promoters. Nonetheless, unlike pre-propagated virus inoculums, all Pol II transcripts would have experienced a single round of error introduction. As a result, this class of errors was easily accounted for by including the RTRC control.

### Mutation spectrum of TCVdMP_sg2R (-) strands

Aside from the 11 G-to-A errors discussed above, the remaining 12 errors specific to TCVdMP_sg2R (-) strands encompassed 8 different classes (Table 3). While the 3 remaining classes of transitions (A-to-G, U-to-C, C-to-U) were all detected twice, only one transversion class (C-to-A) was detected twice. By contrast, three transversions, U-to-A, G-to-C, and U-to-G, were completely absent. Intriguingly, despite the relative rarity of transversions, three of them, A-to-U, C-to-G, and A-to-C, were actually found to co-exist in the same cDNA clone (* in Table 3), along with another mutation that was a G-to-A transition [Fig 2. All 4 mutations are highlighted with light blue boxes. Also note that here the error identities were converted to their (+)-strand complements]. The simultaneous occurrence of these four errors in the same clone raised the possibility that one of them might be a primary, fidelity-relaxing error, causing the TCV RdRp to preferentially mis-incorporate transversions. It was further possible that such fidelity-relaxing mutation could have been introduced by Pol II, given its relatively high error rate as discussed above. Consistent with these deliberations, all four of the errors caused amino acid (aa) identity changes in p88 (Fig 2).

### Long reads unveil an error distribution pattern suggestive of one replication cycle per cell

The relatively long inserts (2,363 bp, or 58% of the TCV genome) of our clones permitted us to inspect the pattern of error distribution in the clones in comparison with random error occurrence, with an error rate of 8.47 X 10^−5^. For this we must first adjust the number of errors in TCVdMP_sg2R-derived clones against the error number of RTRC-derived clones. The first adjustment was to subtract the 23 errors with 13 identified in RTRC-derived clones, arriving at an adjusted error number of 10. The second adjustment dealt with the two clones that contained 4 errors each before adjustment. We assumed that they each contained 2 errors that were introduced by Pol II and/or other steps of the experiment. This assumption was based on two rationales: (i) the ratio of RTRC errors vs TCVdMP_sg2R errors was 13/23 = 0.57, meaning that among the 8 errors found in these two clones, 4 – 5 (8 X 0.57) were probably introduced by Pol II or other experimental steps; (ii) two of the RTRC-derived clones contained 2 errors each. As a result of the adjustments, the number of clones with 1 and 2 errors became 6 and 2, respectively, adding to a total of 10 errors (Table 4, Adjusted column). The remaining 42 clones would be error-free.

**Table 4.** Poisson Distribution prediction of probabilities (p) of clones containing 0, 1, 2, 3, and 4 errors (k), and the deduced numbers of clones out of 50 sequenced (#/50). The observed numbers of clones, adjusted or raw, are listed for comparison. Calculations were carried out for two error frequencies (λ): 0.2 (adjusted) and 0.46 (unadjusted), per 2,363-nt (-) strand fragment.

| k | $\lambda = 0.20$ | | | $\lambda = 0.46$ | | |
| --- | --- | --- | --- | --- | --- | --- |
|  | p | #/50 | Adjusted | p | #/50 | Raw |
| 0 | 0.819 | 41 | 42 | 0.631 | 31.5 | 33 |
| 1 | 0.164 | 8.2 | 6 | 0.290 | 14.5 | 15 |
| 2 | 0.016 | 0.8 | 2 | 0.067 | 3.3 | 0 |
| 3 | 0.001 | 0 | 0 | 0.010 | 0.5 | 0 |
| 4 | 0 | 0 | 0 | 0.001 | 0 | 2 |

In comparison, an error rate of 8.47 X 10^−5^, assuming a single cycle of replication, would predict that an average 2,363-nt fragment had a chance of 0.2 (8.47 X 10^−5^ X 2,363) to contain one error. Nevertheless, due to the stochastic nature of error occurrence, the probability exists that some of the (-) strand fragments might contain 2 or even more errors. Such probabilities can be calculated using the Poisson distribution formula below, with p representing the probability of having *k* number of errors occurring in a fragment, given an error frequency of λ per fragment (0.20 in our case):

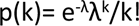

The probabilities of having 0, 1, 2, 3, and 4 errors, along with the predicted numbers of clones (out of a total of 50) under each category, are listed in Table 4. The fact that the predicted numbers closely tracked the adjusted numbers for the 0, 1, 3, and 4 error categories indicated that most of the (-) strands profiled were direct progeny of the primary replication-initiating (+) strands. Put it differently, the progeny (+) strands rarely initiated new replication in the cells of their own genesis. Given the slightly-more-than-expected occurrence of clones with two errors (2 versus 0.8), one might argue that a fraction of progeny genomes did re-initiate replication without exiting parental cells. While such events could not be completely ruled out, they are unlikely to be frequent.

To lay out the rationales for this argument, we first imagine that the newly synthesized progeny genomes repeat replication for just one cycle. Note that only one copy of Pol II-transcribed TCV genome could initiate viral replication in each agro-infiltrated *N. benthamiana* cells to synthesize viral (-) strands (26, 28, 29). As a result, second-cycle replication by even a small fraction of progeny (+) strands would produce enough secondary (-) strands to make themselves detectable. Such secondary (-) strands would have gone through two more copying steps: (-) to (+), then back to (-), meaning they would be expected to contain 3 times as many errors as primary (-) strands. Further extrapolating this thought experiment for two more replication cycles, we could see that (-) strands of 3^rd^ and 4^th^ generations would have 5 and 7 times as many errors as the primary (-) strands. Therefore, if TCV replication were to repeat for multiple cycles per cell, we would detect (-) strands that contain errors at varying frequencies, with the overall pattern deviating dramatically from Poisson distribution. Thus, the fact that the observed pattern conformed to Poisson distribution was highly consistent with the stamping machine replication mode of one cycle per cell.

Similar conclusions could also be drawn using the raw data without adjustment (Table 2 and Table 4). Although Pol II errors occupied a substantial fraction of raw errors, these errors should maintain a constant frequency irrespective of the number of replication cycles, as Pol II transcripts could not re-enter transcription/replication independent of viral replication. Put differently, Pol II errors were introduced through one single round of transcription at a relatively constant rate (approximately 9.31 X 10^−5^) that was very similar to TCV RdRp (8.47 X 10^−5^). As a result, the raw data with a total number of errors at 23 for TCVdMP_sg2R (-) strands could also be treated as if they replicated with a higher error frequency (λ) of 0.46 per 2,363-nt fragment. Repeating the Poisson Distribution calculation, the predicted numbers of clones with 0, 1, 2, 3, and 4 errors would be 31.5, 14.5, 3.3, 0.5, and 0 (Table 4). Again the clone numbers with 0, 1, and 3 errors closely matched Poisson Distribution. By contrast, the clone numbers with 2 and 4 errors moderately deviated from Poisson distribution predictions. These results reinforced the conclusion that most of the (-) strands were products of one-cycle replication. That 2 clones had 4 errors may indicate a small fraction of (-) strands arose from more than one replication cycles, or from more frequent error introduction by RdRp variants with primary, fidelity-relaxing mutation(s).

It is important to note that rare multi-cycle replication events could themselves be due to mutations that perturbed the genetic control of replication cycles. If one-cycle replication is an evolutionarily selected, genome-encoded trait of TCV, one can expect this trait being occasionally undermined by mutations in the gene(s) encoding this trait. Rare viral mutants containing such mutations, before being purged by natural selection, can amplify themselves to high numbers through multi-cycle replication, and concomitantly incurring additional errors, hence becoming detected by error-profiling. Similarly, since the error rate of a viral RdRp is also optimized through natural selection, it too could be relaxed by occasional mutations. Such occasional error rate relaxation would be expected to introduce more errors in the progeny genomes, thereby interfering with both the estimation of error rates and determination of per-cell replication cycles. In short, while we cannot completely rule out occasional second-cycle replication, such occurrence could very well reflect primary mutation events weakening other genetically encoded traits.

We hasten to add two qualifications to the one-replication-cycle-per-cell conclusion. First, one replication cycle per cell should not be taken to mean just one (-)-strand copy from every (+)-strand template. Rather, we argue that each of the (+) strands templated the synthesis of numerous (-)-strand copies, and each of the (-) strands then templated the synthesis of numerous (+) strands. This expands the stamping machine model proposed by others by proposing that “stamping” can occur on both founding (+) strands and their (-)-strand intermediates (13, 34). Second, sgRNAs produced by TCV and other similar viruses need not to obey the one cycle rule imposed on genomic RNAs. This is because sgRNA production is known to be dependent on active replication, hence sgRNA (-) strands must be copied from progeny (+) RNAs. However, sgRNAs are not inherited by the next generation of viruses, thus can afford to harbor more errors.

It should be noted that at least two other RNA viruses, namely bacteriophage ɸ6 and turnip mosaic virus, have been reported to replicate through a predominantly stamping machine mode (13, 34). However, this mode does not appear to apply to the replication of poliovirus (PV), because “on average the (PV) viral progeny produced from each cell are approximately five generations removed from the infecting virus” (35). This is despite the fact that the error rate of PV RdRp, at 9.0 X 10^−5^ (19), is strikingly similar to that of TCV RdRp. The simplest explanation appears to be that TCV and PV are different viruses that infect drastically different host cells. For example, TCV synthesizes some of the viral proteins (p8, p9, p38) using subgenomic mRNAs produced in the infected cells, whereas PV synthesizes all viral proteins in the form of a continuous polyprotein precursor that is subsequently proteolytically processed to yield mature proteins. As a result, there exists a possibility that some, or even most, of PV RNAs in the infected cells are dedicated mRNAs that do not become part of progeny viruses, thus could afford to tolerate more errors. Aside from this, it is also possible that the sequence reads profiled by the PV study (35) were too short to permit the differentiation of primary errors inherent of the wildtype RdRp from secondary errors introduced by mutated forms of RdRp that were more error-prone. Finally, errors pre-existing in the PV inoculums might have contributed higher error frequencies in the PV study. Careful assessment of these possibilities should reveal the underlying evolutionary rationales for the different replication modes.

### Two lethal TCV mutants captured through (-)-strand profiling

To determine whether the TCVdMP_sg2R-specific errors would affect the functionality of the p28/p88 proteins, the (-)-strand errors were converted to their (+)-strand complements and assessed for their potential to alter the identity of p28/p88 aa residues. Six of the 23 errors (C399U, C627U, C972U, C1290U, A1539G, U2182C) were predicted to maintain the aa residues of wildtype RdRp (Fig 2, middle, unaltered aa are underlined). Most of the remaining errors were predicted to cause aa changes that did not appear to be seriously debilitating (e.g. A113V, A133V, T172M, A233V, G547S, E748Q), though their specific impacts remain to be investigated. However, several mutations did have the potential to cause more drastic aa changes, including S153L, R204H, R273C, Y677N, F726L, and L740R. Finally, the mutation G2332U caused the most dramatic aa change by converting G757 into a premature stop codon (Fig 2, middle, red font). Interestingly, this mutation was part of the second clone with 4 errors (Fig 2, middle, orange boxes), suggesting that this mutation was likewise among the secondary errors caused by a more error-prone mutant RdRp.

We next wondered if TCV mutants containing the two sets of 4 errors were still capable of productive replication. To this end, we incorporated the two sets of errors into TCVdMP_sg2R backbone, creating mut17 (C401U/G674A/C972U/G2332U) and mutY6 (C521U/Y677N/U2282G/G2305C), respectively. Both mutants were brought into *N. benthamiana* leaf cells via agro-infiltration, and their replication levels assessed with Northern blotting. As shown in Fig 3, Neither of the mutants accumulated TCV RNAs to detectable levels. Therefore, although mutants with multiple errors could emerge in single cell infections, most of them would have lost the ability to replicate in single cells, thus likely escaping detection by other error-profiling techniques. We are currently examining the individual errors making up of these two mutants to determine whether any of them lowers the RdRp fidelity or permits more replication cycles per cell. Such follow-up examination is expected to reveal additional insights on the evolutionary mechanisms of (+) RNA virus replication.

**Figure 3.**
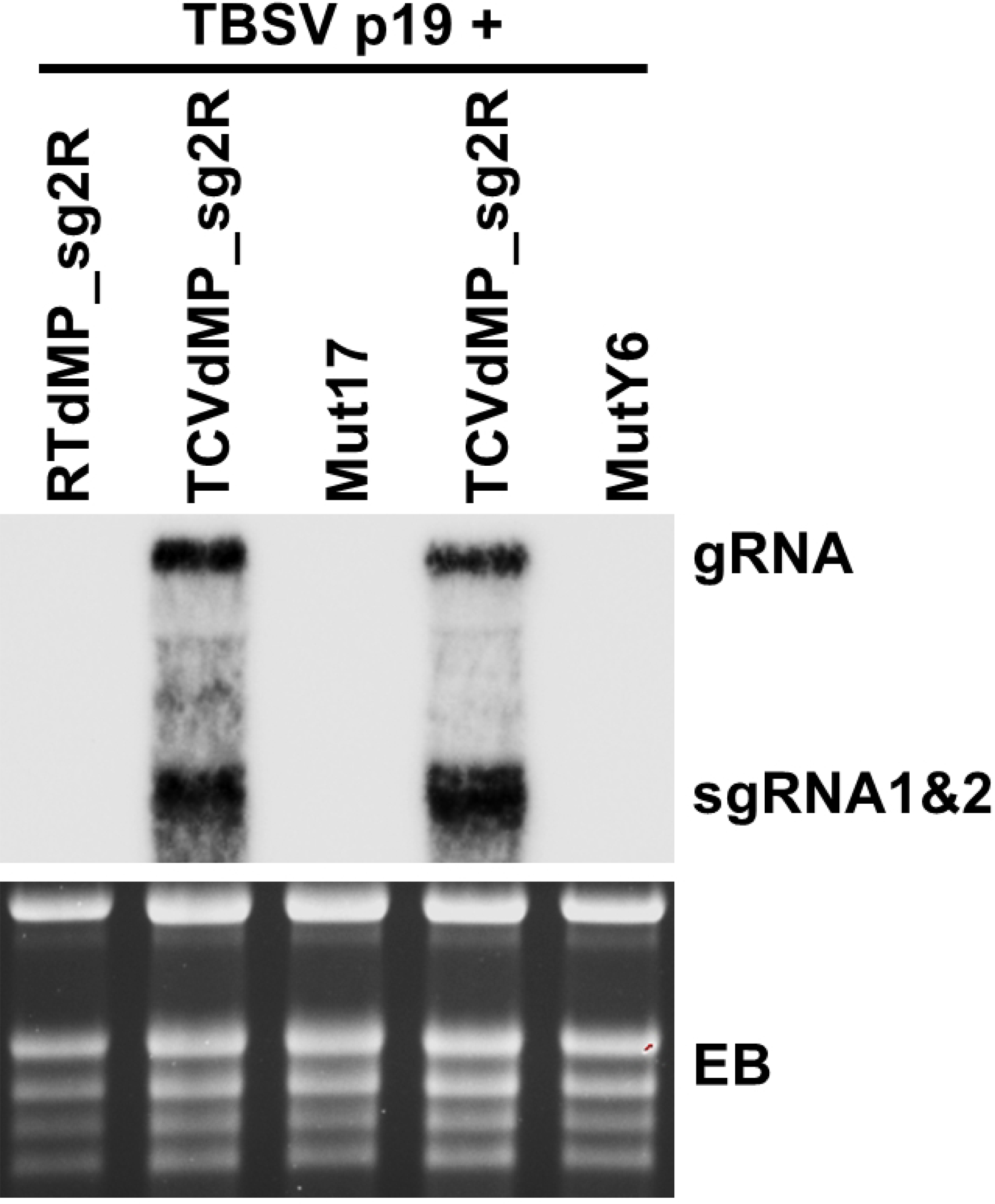
Inability of mut17 and mutY6 to replicate in *N. benthamiana* cells. Northern blotting was carried out using total RNA samples extracted from *N. benthamiana* leaves agro-infiltrated with the constructs indicated on the top, with TCV-specific oligonucleotide probes (sequences available upon request). EB: ethidium bromide-stained gel showing equal loading.

## Conclusions

Here we report an effort to obtain a more accurate estimation of the error rate of a (+) RNA virus. Our effort focused exclusively on errors incurred in (-)-strand replication intermediates produced in single cells in which replication was launched with transcribable viral cDNA. We additionally included controls that transcribed only (+) or (-) strands, ensuring exclusive capture of replication-generated (-) strands, and meaningful estimation of experiment-borne errors. These efforts allowed us to compute an error rate for TCV RdRp that was no higher than 1.947 X 10^−4^. Further analyses revealed that the host cell Pol II could have contributed errors at up to 9.31 X 10^−5^. Factoring potential errors introduced by other experimental steps, we arrived at a TCV RdRp error rate of 8.47 X 10^−5^ s/n/c. Moreover, purposeful production of long continuous cDNA fragments (2,363 bp) permitted us to map errors to their respective clones with high confidence, revealing an error distribution pattern consistent with the replication mode of one cycle per cell. Finally, our new procedure also permitted the capture of lethal errors that would have missed with some of the earlier procedures. Our findings offer novel insights on the mechanism of (+) RNA virus replication.

## Acknowledgements

We thank the USDA ARS Maize and Soybean viruses Group for generous equipment sharing. Members of the Qu lab are greatly appreciated for discussions and technical assistances. This study is supported in part by the NSF grant 1758912. CPC is supported in part by a scholarship from the Coordenação de Aperfeiçoamento de Pessoal de Nível Superior - Brasil (CAPES) – Finance Code 001. SM is supported in part by Japan Society for the Promotion of Science (JSPS) KAKENHI grant 21K05591.

## Materials and Methods

### Constructs

Both TCVdMP_sg2R and RTdMP_sg2R are binary constructs modified from TCV_sg2R and RT_sg2R (21, 36), respectively, by creating a 92-nt deletion (positions 2,425-2,516) within the p8/p9 coding region. This deletion was previously shown to abolish TCV cell-to-cell movement without affecting the production of sgRNAs (14). Note that the deletion is outside the region being subject to RT-PCR and error profiling (see later). The insert of the construct RTRC consists of the first 2,489 nt of RTdMP_sg2R, in reverse-complemented orientation, cloned immediately downstream of P35S in pAI101 (22, 27).

### Agrobacterium infiltration (agro-infiltration)

All DNA constructs destined for testing in *N. benthamiana* plants were transformed into electrocompetent A. tumefaciens strain C58C1 via electroporation using the AGR setting of a Bio-Rad Micropulser Electroporator. Briefly, 5 µl of the plasmid DNA was mixed with 40 µl of agro cells and maintained on ice until electroporation. After electroporation, 900 µl of SOB media was added and the suspension was incubated at 28 °C for one hour. Selection was carried out on solid Terrific Broth (TB) media containing rifampicin, gentamycin, and kanamycin. Successful introduction of the plasmid was confirmed using colony PCR. A single colony confirmed to have the desired plasmid was used to inoculate 3 ml TB liquid media with the same antibiotics, and incubated overnight in a 28 °C shaker (220 rpm). The culture was diluted 1:100 with fresh TB liquid media and incubated under the same conditions for another night. The second culture was centrifuged at 4,000 rpm for 20 min, and resuspended in agroinfiltration buffer (10 mM MgCl2, 10 mM MES, and 100 µM acetosyringone). All suspensions were diluted to OD600 = 1 and incubated at room temperature for 3 hours. They were then mixed and introduced into leaves of young *N. bethamiana* plants via a small wound, using a needleless syringe.

### Confocal microscopy

Four days after agro-infiltration, leaf discs were collected from the plants. Confocal microscopy was performed at the Molecular and Cellular Imaging Center (MCIC), the Ohio Agricultural Research and Development Center, using a Leica DMI6000 laser confocal scanning microscope. To detect GFP and mCherry fluorescence, sequential excitation at 488 nm and 587 nm was provided by argon and helium-neon 543 lasers, respectively.

### Strand-specific reverse transcription-polymerase chain reaction (RT-PCR), cloning of the PCR products

To generate the (-)-strand-specific cDNA of TCV, we adopted the procedure reported by Plaskon and colleagues (33). Specifically, to initiate (-)-strand-specific RT, we used the primer ssRT-Tg3-TCV25F (5’-TTGTAAAACGACGGCCAGT <u>GAGCT</u> <u>C</u>GCCTAAAATTGCCCTCA-3’). This primer comprised three sections: a 19-nt non-TCV tail at the 5’ end, derived from the sequence of pUC19, was followed by a 5-nt linker (GAGCT) in the middle, and an 18-nt 3’ terminus derived from positions 25-42 of TCV (+) strand. The GAGCT linker along with the C downstream created a SacI site (underlined). The RT was carried out with the RevertAid Reverse Transcriptase (Thermo Scientific) following the Manufacturer’s instructions. For PCR, we used the primer ssRT-Tg3 (500 nM), whose sequence was identical to the first 22 nt of ssRT-Tg3-TCV25F, to pair with a primer mix consisting of ssRT-Tg4-TCV2425R (25 nM) and ssRT-Tg4 (500 nM). The sequence of ssRT-Tg4-TCV2425R (5’-CTATGACCATGATTACGCC<u>AAGCTT</u> CCTTTCTTCCGTTTTCCTGT-3’) consisted of a 5’ 25-nt tail derived from pUC19, and a 20-nt 3’ portion complementary to positions 2406-2425 of TCV (+) strand; whereas that of ssRT-Tg4 corresponded to the first 22 nt of ssRT-Tg4-TCV2425R. PCR was carried out with the Phusion High Fidelity Master Mix (Thermo Scientific), according to Manufacturer’s instructions.

The PCR-amplified 2,450-bp cDNA fragment was gel-purified, and cloned into pUC19 (digested with EcoRI plus HindIII) using the NEBuilder kit (New England Biolabs). The cloning products were then transformed into E. coli. Plasmids isolated from the E. coli colonies were digested with SacI plus HindIII to verify the size of inserts. Those with inserts of expected size were subject to Sanger sequencing with two sets of primers. The four primers used for sequencing earlier batches of plasmids were: TCV-755R, -503F, -1104F, and -1711F. The three primers used for sequencing the later batches of plasmids were: TCV-946R, -832F, and -1627F. Sequences of these primers are available upon request.

### Northern blotting

Total RNA was extracted from agro-infiltrated *N. benthamiana* leaves using the Direct-zol RNA Miniprep kit (Zymo Research, Irvine, CA). To ensure consistency, six equivalent leaf sections derived from infiltrated leaves of three different plants were pooled before RNA extraction. The RNA extraction procedure included a DNase treatment step that removed DNA contamination. The RNA was then quantified with NanoDrop and subjected to Northern blotting as described (21, 22).

## Notes

### Competing Interest Statement

The authors have declared no competing interest.

## References

1. Duffy S. 2018. Why are RNA virus mutation rates so damn high? PLoS Biol 16:e3000003–e3000003.

2. Peck KM, Lauring AS. 2018. Complexities of Viral Mutation Rates. J Virol 92:e01031–17.

3. Domingo E, García-Crespo C, Lobo-Vega R, Perales C. 2021. Mutation Rates, Mutation Frequencies, and Proofreading-Repair Activities in RNA Virus Genetics. Viruses 13:1882.

4. Elena SF, Sanjuán R. 2005. Adaptive value of high mutation rates of RNA viruses: separating causes from consequences. J Virol 79:11555–11558.

5. Malpica JM, Fraile A, Moreno I, Obies CI, Drake JW, García-Arenal F. 2002. The Rate and Character of Spontaneous Mutation in an RNA Virus. Genetics 162:1505.

6. Sanjuán R, Agudelo-Romero P, Elena SF. 2009. Upper-limit mutation rate estimation for a plant RNA virus. Biol Lett 5:394–396.

7. Tromas N, Elena SF. 2010. The rate and spectrum of spontaneous mutations in a plant RNA virus. Genetics, 2010/05/03 ed. 185:983–989.

8. Lafforgue G, Martínez F, Sardanyés J, de la Iglesia F, Niu Q-W, Lin S-S, Solé RV, Chua N-H, Daròs J-A, Elena SF. 2011. Tempo and Mode of Plant RNA Virus Escape from RNA Interference-Mediated Resistance. J Virol 85:9686–9695.

9. Acevedo A, Brodsky L, Andino R. 2014. Mutational and fitness landscapes of an RNA virus revealed through population sequencing. Nature 505:686–690.

10. Geller R, Estada Ú, Peris JB, Andreu I, Bou J-V, Garijo R, Cuevas JM, Sabariegos R, Mas A, Sanjuán R. 2016. Highly heterogeneous mutation rates in the hepatitis C virus genome. Nat Microbiol 1:16045.

11. de la Iglesia F, Martínez F, Hillung J, Cuevas JM, Gerrish PJ, Daròs J-A, Elena SF. 2012. Luria-Delbrück Estimation of Turnip Mosaic Virus Mutation Rate *In Vivo*. J Virol 86:3386–3388.

12. Pauly MD, Procario MC, Lauring AS. 2017. A novel twelve class fluctuation test reveals higher than expected mutation rates for influenza A viruses. eLife 6:e26437.

13. Chao L, Rang CU, Wong LE. 2002. Distribution of Spontaneous Mutants and Inferences about the Replication Mode of the RNA Bacteriophage ␾6. J VIROL 76:3276–3281.

14. Li W, Qu F, Morris TJ. 1998. Cell-to-Cell Movement of Turnip Crinkle Virus Is Controlled by Two Small Open Reading Frames That Functionin trans. Virology 244:405–416.

15. Lucas WJ. 2006. Plant viral movement proteins: Agents for cell-to-cell trafficking of viral genomes. Virol 50th Anniv Issue 344:169–184.

16. Qu F, Ren T, Morris TJ. 2003. The Coat Protein of Turnip Crinkle Virus Suppresses Posttranscriptional Gene Silencing at an Early Initiation Step. J Virol 77:511–522.

17. Cao M, Ye X, Willie K, Lin J, Zhang X, Redinbaugh MG, Simon AE, Morris TJ, Qu F. 2010. The Capsid Protein of Turnip Crinkle Virus Overcomes Two Separate Defense Barriers To Facilitate Systemic Movement of the Virus in Arabidopsis. J Virol 84:7793–7802.

18. Qu F, Ye X, Morris TJ. 2008. Arabidopsis DRB4, AGO1, AGO7, and RDR6 participate in a DCL4-initiated antiviral RNA silencing pathway negatively regulated by DCL1. Proc Natl Acad Sci 105:14732–14737.

19. Sanjuán R, Domingo-Calap P. 2016. Mechanisms of viral mutation. Cell Mol Life Sci 73:4433–4448.

20. Lin J, Guo J, Finer J, Dorrance AE, Redinbaugh MG, Qu F. 2014. The Bean Pod Mottle Virus RNA2-Encoded 58-Kilodalton Protein P58 Is Required in *cis* for RNA2 Accumulation. J Virol 88:3213–3222.

21. Zhang S, Sun R, Perdoncini Carvalho C, Han J, Zheng L, Qu F. 2021. Replication-Dependent Biogenesis of Turnip Crinkle Virus Long Noncoding RNAs. J Virol 95:e00169–21.

22. Sun R, Zhang S, Zheng L, Qu F. 2020. Translation-Independent Roles of RNA Secondary Structures within the Replication Protein Coding Region of Turnip Crinkle Virus. Viruses 12:350.

23. Gout J-F, Li W, Fritsch C, Li A, Haroon S, Singh L, Hua D, Fazelinia H, Smith Z, Seeholzer S, Thomas K, Lynch M, Vermulst M. 2017. The landscape of transcription errors in eukaryotic cells. Sci Adv 3:e1701484.

24. Gout J-F, Thomas WK, Smith Z, Okamoto K, Lynch M. 2013. Large-scale detection of in vivo transcription errors. Proc Natl Acad Sci 110:18584–18589.

25. Qu F, Morris TJ. 2002. Efficient Infection of Nicotiana benthamiana by Tomato bushy stunt virus Is Facilitated by the Coat Protein and Maintained by p19 Through Suppression of Gene Silencing. Mol Plant-Microbe Interactions® 15:193–202.

26. Zhang X-F, Sun R, Guo Q, Zhang S, Meulia T, Halfmann R, Li D, Qu F. 2017. A self-perpetuating repressive state of a viral replication protein blocks superinfection by the same virus. PLOS Pathog 13:e1006253.

27. Guo Q, Zhang S, Sun R, Yao X, Zhang X-F, Tatineni S, Meulia T, Qu F. 2020. Superinfection Exclusion by p28 of Turnip Crinkle Virus Is Separable from Its Replication Function. Mol Plant Microbe Interact 33:364–375.

28. Qu F, Zheng L, Zhang S, Sun R, Slot J, Miyashita S. 2020. Bottleneck, Isolate, Amplify, Select (BIAS) as a mechanistic framework for intracellular population dynamics of positive-sense RNA viruses. Virus Evol 6.

29. Perdoncini Carvalho C, Ren R, Han J, Qu F. 2022. Natural Selection, Intracellular Bottlenecks of Virus Populations, and Viral Superinfection Exclusion. Annu Rev Virol 9:4.1–4.17.

30. Qu F, Ye X, Hou G, Sato S, Clemente TE, Morris TJ. 2005. RDR6 Has a Broad-Spectrum but Temperature-Dependent Antiviral Defense Role in Nicotiana benthamiana. J Virol 79:15209– 15217.

31. Garcia-Ruiz H, Takeda A, Chapman EJ, Sullivan CM, Fahlgren N, Brempelis KJ, Carrington JC. 2010. Arabidopsis RNA-Dependent RNA Polymerases and Dicer-Like Proteins in Antiviral Defense and Small Interfering RNA Biogenesis during Turnip Mosaic Virus Infection. Plant Cell 22:481–496.

32. Wang X-B, Wu Q, Ito T, Cillo F, Li W-X, Chen X, Yu J-L, Ding S-W. 2010. RNAi-mediated viral immunity requires amplification of virus-derived siRNAs in Arabidopsis thaliana. Proc Natl Acad Sci 107:484–489.

33. Plaskon NE, Adelman ZN, Myles KM. 2009. Accurate Strand-Specific Quantification of Viral RNA. PLOS ONE 4:e7468.

34. Martinez F, Sardanyes J, Elena SF, Daros J-A. 2011. Dynamics of a Plant RNA Virus Intracellular Accumulation: Stamping Machine vs. Geometric Replication. Genetics 188:637–646.

35. Schulte MB, Draghi JA, Plotkin JB, Andino R. 2015. Experimentally guided models reveal replication principles that shape the mutation distribution of RNA viruses. Elife 4:e03753.

36. Zhang S, Sun R, Guo Q, Zhang X-F, Qu F. 2019. Repression of turnip crinkle virus replication by its replication protein p88. Virology 526:165–172.

